# Antioxidant Activity via Free Radical Scavenging of Pitavastatin and Its Hydroxylated Metabolites. A Quantum Chemical Attempt Aiming to Assist Drug Development

**DOI:** 10.1101/2023.04.12.536546

**Authors:** Ioan Bâldea

**Affiliations:** Theoretical Chemistry, Heidelberg University, Im Neuenheimer Feld 229, D-69120 Heidelberg, Germany

**Keywords:** free radical scavenging activity, antioxidant mechanisms, direct hydrogen atom transfer (HAT), stepwise electron transfer proton transfer (SETPT), sequential proton loss electron transfer (BDE), thermochemstry, quantum chemistry

## Abstract

Statins form a class of drugs often administered in a variety of cardiovascular diseases, for which their antioxidant capacity appears particularly relevant. Although experiments have long provided empirical evidence that statins can suppress various oxidation pathways, theoretical attempts to quantify the antioxidant activity of statins (read, atorvastatin ATV, because this is the only one studied so far) were not published until last year. Molecular and clinical differences of stains trace back to the ring attached to the statin’s active moiety. This can be, e.g., a pyrrole, as the case of the aforementioned ATV or a quinoline, as the case of pitavastatin (PVT), which represents the focus of the present work. Extensive results reported here for PVT and derivative include the thermodynamic antioxidant descriptors (bond dissociation enthalpy BDE, adiabatic ionization potential IP, proton dissociation enthalpy PDE, proton affinity PA, and electron transfer enthalpy ETE) related to the three antioxidant mechanisms (hydrogen atom transfer HAT, stepwise electron transfer proton transfer SETPT, sequential proton loss electron transfer SPLET). Our particular emphasis is on the PVT’s hydroxylated derivatives wherein a hydroxy group replaces a hydrogen atom either on the quinoline core (Q-hydroxylated metabolites) or on the fluorophenyl ring (F-hydroxylated metabolites). Our calculations indicate that both the Q- and F-hydroxylated metabolites possess antioxidant properties superior to the parent PVT molecule. Given the fact that, to the best of our knowledge, no experimental data for the antioxidant potency of PVT and its hydroxylated derivatives exist, this is a theoretical prediction, and we Given the fact that, to the best of our knowledge, no experimental data for the antioxidant potency of PVT and its hydroxylated derivatives exist, this is a theoretical prediction for the validation of which we aim hereby to stimulate companion experimental in vivo and in vitro investigations and inspire pharmacologists in further drug developments.

## 1. Introduction

Cardiovascular disease (CVD)— which includes, e.g., heart attack, atherosclerosis, angina, peripheral artery disease, and stroke— is sadly reputed to cause about one third of all deaths worldwide [1], and blood low-density lipoprotein cholesterol concentration is recognized as a major CVD risk factor [2,3].

Cholesterol is the final product of the mevalonate pathway. Therefore, in cholesterol biosynthesis, the mevalonate pathway is the metabolic pathway of paramount importance. The first step in cholesterol biosynthesis is the production of 3-hydroxy-3-methyl-glutaryl coenzyme A (or, alternatively, *β* -hydroxy *β* -methylglutaryl coenzyme, HMG-CoA), which is eventually converted to mevalonate. HMG-CoA conversion to mevalonate is catalyzed by the enzyme HMG-CoA reductase, whose level is regulated through a multivalent feedback mechanism [4]. This is the limiting rate reaction in cholesterol biosynthesis [5].

Statins are a class of drugs that act as powerful inhibitors of the mevalonate biosynthetic pathway. Strongly inhibiting HMG-CoA reductase [6–10], they prevent cholesterol synthesis. Due to their well-proven efficiency in reducing blood low-density lipoprotein cholesterol (LDL-C) concentration, statins are widely prescribed medications for both primary and secondary CVD prevention [11–21].

Low-density lipoprotein cholesterol triggers oxidative stress [22–24]. The oxidative stress is an extremely dangerous phenomenon caused by the rapid production of free radicals in human body [25–29]. Free radicals can seriously damage a variety of biomolecules, generating thereby numerous pathological processes that include CVD [30–41]. Antioxidants are molecules that can scavenge free radicals by hydrogen atom donation. Statins are known to contribute to oxidative stress reduction [42]. From this perspective, quantifying statins’ antioxidant capacity and their related free radical scavenging activity [42–44], is an aspect that certainly deserves consideration.

Although numerous experimental studies drew attention on the statins’ antioxidant ability to act as efficient free radical scavengers [42–51], theoretical investigations of these topics relying on quantum chemical calculations were not published until last year [52–55]. Encouragingly, efforts in this direction turned out to be rewarding. Noteworthily, they succeeded to explain [53] the long standing conundrum regarding the reason why ortho- and para-hydroxylated atorvastatin metabolites can scavenge free radicals that the parent atorvastatin cannot [47].

In chemistry, hydroxylation is a common process that introduces a hydroxy group (– OH) into an organic compound. Putting in a more medical context, introduction of hydroxy group into a drug, often occurring by replacing a hydrogen atom on an aromatic ring, is the most common reaction type in phase I metabolism. It typically produces a chemically stable and more polar hydroxylated metabolite than the parent drug [56]. Hydroxylation can create additional functional sites available, enabling thereby adequate tuning of the chemical reactivity.

Because in vivo atorvastatin is metabolized to ortho- and para-hydroxy metabolites, the afore-mentioned finding of ref. [53] is particularly important. As a matter of fact, in more than two third of cases the inhibition of the HMG-CoA reductase was attributed to active hydroxylated metabolites rather than to the parent non-hydroxylated species [47,52,57–59].

The foregoing presentation made it clear why investigations of the antioxidant potency of other widely administered statins as well as their hydroxylated metabolites are highly desirable. As a further step undertaken in this direction, in this paper we will focus on the antioxidant properties of pitavastatin (PVT) [9,42,60,61,61–64] and its hydroxylated derivatives. To legitimate our present choice (i.e., PVT), we refer to previous reports presenting empirical evidence that treatment with pitavastatin could decrease oxidative stress (see [42] and citations therein).

PVT (Figure 1) is a synthetic statin patented in 1987 and approved for medical use in 2003 [65]. PVT is among the most potent low-density lipoprotein (LDL) cholesterol-lowering statins. It is well tolerated (low incidence of adverse events), is rapidly absorbed, and has the highest (*∼* 80%) overall bioavailability of any statin.

**Figure 1.**
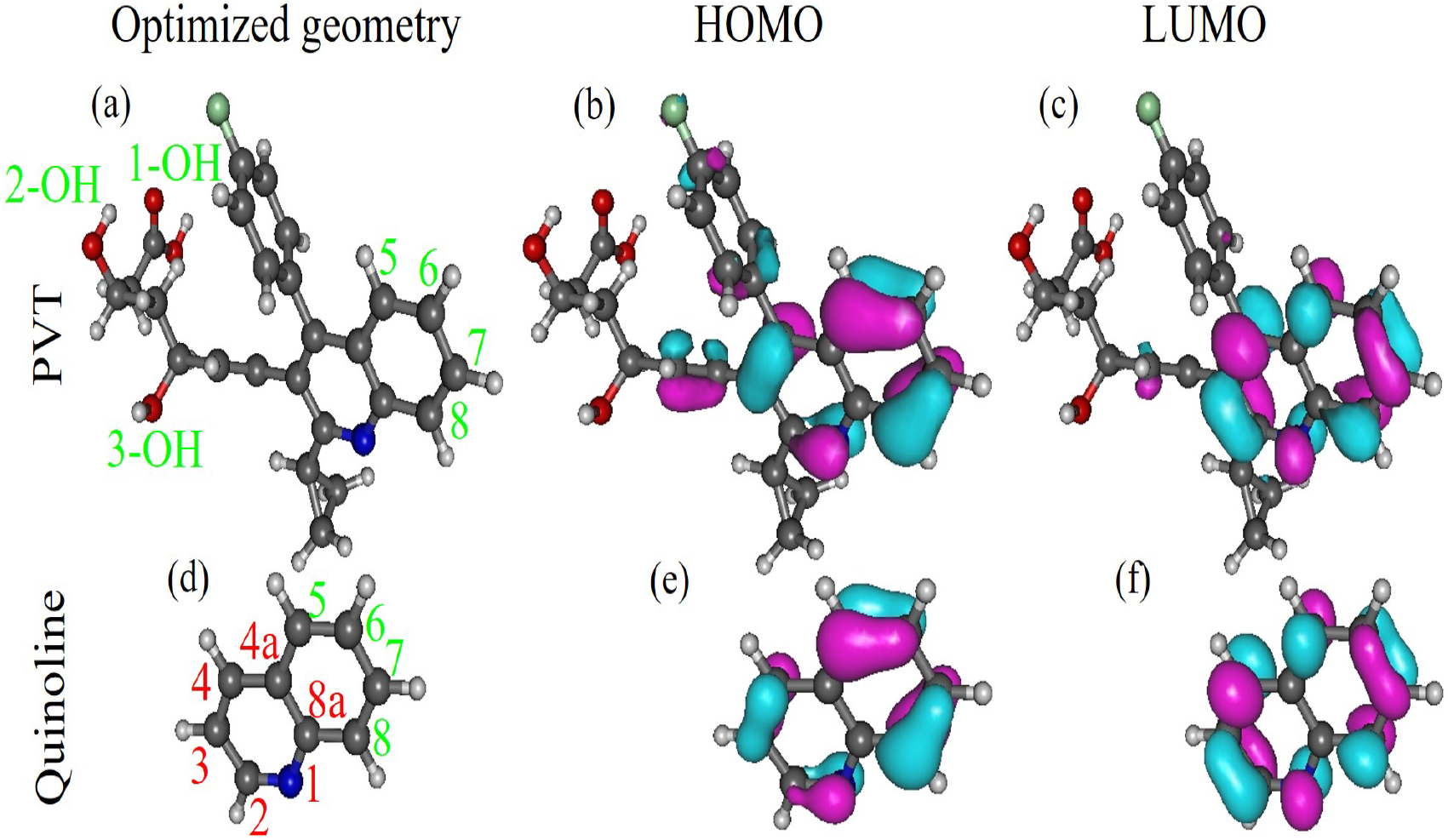
Pitavastatin contains a trisubstituted quinoline core and differs from other synthetic statins by incorporating at position 2 a cyclo-propyl moiety rather than the more typical iso-propyl ring substituent contained by other synthetic statins [60]. Figures generated using Gabedit 2.5.1 [92].

## 2. Computational details

The quantum chemical calculations for presently considered PVT and its hydroxylated derivatives were done using the GAUSSIAN 16 suite of programs [66]. Because they are similar to those reported in our recent studies (e.g., [53–55], only a few details will be given below. Geometry optimizations without constraints, frequency calculations (checking that all vibrational frequencies were real), and electronic energies were done at the DFT level of theory. For consistency with our recent and ongoing work we used the hybrid B3LYP exchange correlation functional [67–70] and 6-31+G(d,p) basis sets [71,72].

Similar to other cases studied recently [73–76], spin contamination did not appear to be an issue for the unrestricted spin (UB3LYP) approaches, as witnessed by the values of the total spin values ⟨ *S*^2^⟩ before and after annihilation of the first spin contaminant, which were found to be very close to each other. The small differences between the unrestricted and restricted open shell methods revealed that dynamical electron correlations brought about by spin polarization effects are weak; still, as emphasized earlier [75,76], they should make it clear that claims (often made in the literature on antioxidation) on chemical accuracy (*∼* 1 kcal/mol) are totally unrealistic. Achieving chemical accuracy for bond dissociation enthalpies (BDE) and proton affinity (PA)— which are quantities entering the discussion that follows— is often illusory even for extremely computationally demanding state-of-the-art compound model chemistries (see [66]) and much smaller molecular sizes [77].

The solvent was treated within the polarized continuum model (PCM) [78] using the integral equation formalism (IEF) [79]. Choosing methanol as solvent is motivated by the fact that antioxidant assays are very often done in methanolic phase [52,80].

The values of all enthalpies of reaction reported below were computed at the temperature *T* = 298.15 K.

## 3. Results and Discussion

### 3.1. Enthalpies of Reaction Characterizing the Antioxidant Activity

Before presenting results quantifying the antioxidant activity of PVT and its hydroxylated derivatives, in order to make the paper self-consistent, we first provide the reader with the most relevant details on the antioxidant activity via free radical scavenging.

An antioxidant (AXH) can transfer a hydrogen atom (forming a chemical bond X – H with one of its atoms X, e.g., X = O) to a free radical (R^*•*^) via three mechanisms: direct hydrogen atom transfer (HAT), stepwise electron transfer proton transfer (SETPT), and sequential proton loss electron transfer (SPLET). The first is a single-step mechanism, the last two are two-step mechanisms.

Direct hydrogen atom transfer (HAT) is a mechanism involving a single step process [81–83]. For obvious reasons, the pertaining enthalpy of this homolytic bond cleavage process is called bond dissociation enthalpy (BDE)

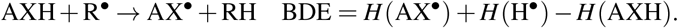

The stepwise electron transfer proton transfer (SETPT) [84–87] is a two step mechanism. In the first step, an electron is transferred (antioxidant’s ionization, and hence the generic term ionization “potential” IP for the corresponding reaction enthalpy). The proton dissociation enthalpy (PDE) characterizes the second step, a reaction involving the transfer of a proton

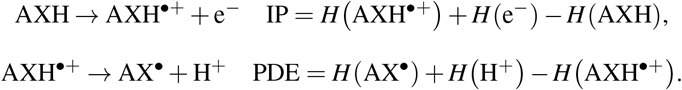

The first step of the sequential proton loss electron transfer (SPLET) mechanism [88,89] is a heterolytic bond cleavage process wherein a proton is transferred from the antioxidant to the free radical. Electron transfer follows in the second step

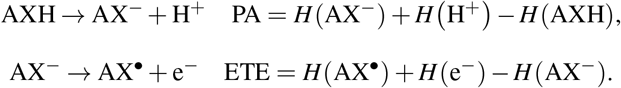

The pertaining reaction enthalpies quantifying these processes are the proton affinity (PA) and the electron transfer enthalpy (ETE), respectively.

In the present case of interest (namely, PVT and its hydroxylated derivatives), X stands for an O atom. In case of other antioxidants, X can also be an N or an S atom. The above definitions should make clear that all aforementioned reaction enthalpies (in particular, IP) are adiabatic rather than vertical properties [90,91]; they should be evaluated at the global electronic energy minima of the various reaction products/reactants.

### 3.2. Characterization of the Pitavastatin (PVT) Molecule

In view of the fact that the present paper is mainly intended to be a study of antioxidant properties rather than a comprehensive quantum chemical study of PVT and related molecular species, we restrict ourselves to merely present the optimized geometries and the spatial distributions of the frontier molecular orbitals (FMO)— highest occupied molecular orbitals (HOMO and lowest unoccupied molecular orbitals (LUMO)— of the relevant molecular species to be analyzed.

Figure 1 depicts the molecular structure of PVT along with that of its trisubstituted quinoline core. The cyclo-propyl moiety attached at quinoline’s position 2 (for positions’ notation, see Figure 1) represents a significant difference from the more typical iso-propyl ring substituent characteristic for other synthetic statins. Figure 1 also depicts the spatial distributions of the frontier (HOMO and LUMO) molecular orbitals. They are interesting because they give a flavor of molecule’s chemical reactivity. By and large, high HOMO densities characterize spatial regions able to donate electrons (molecule’s nucleophilic part) while high LUMO densities correspond to spatial regions favorable to accepting electrons (molecule’s electrophilic part).

Inspection of Figure 1 reveals that the chemical bonds having high HOMO density and the chemical bonds having high LUMO density exhibit complementary spatial location within the PVT molecule. Noteworthily, both these bonds belong to the quinoline core. Comparison with the isolated quinoline molecule reveals that, basically, PVT’s HOMO and LUMO originate from quinoline’s frontier molecular orbitals (HOMO and LUMO). Similar to the isolated quinoline molecule, neither the HOMO nor the LUMO distribution of PVT extends over the (C_4*a*_ – C_8*a*_) carbon-carbon bond where the benzene ring and the pyridine ring are fused together.

PVT can scavenge free radicals by donating hydrogen atoms from the three OH groups it contains (positions 1-OH, 2-OH, and 3-OH in Figure 1). The enthalpies of reaction quantifying PVT’s antioxidant activity are presented in the upper part of Table 1.

**Table 1.**
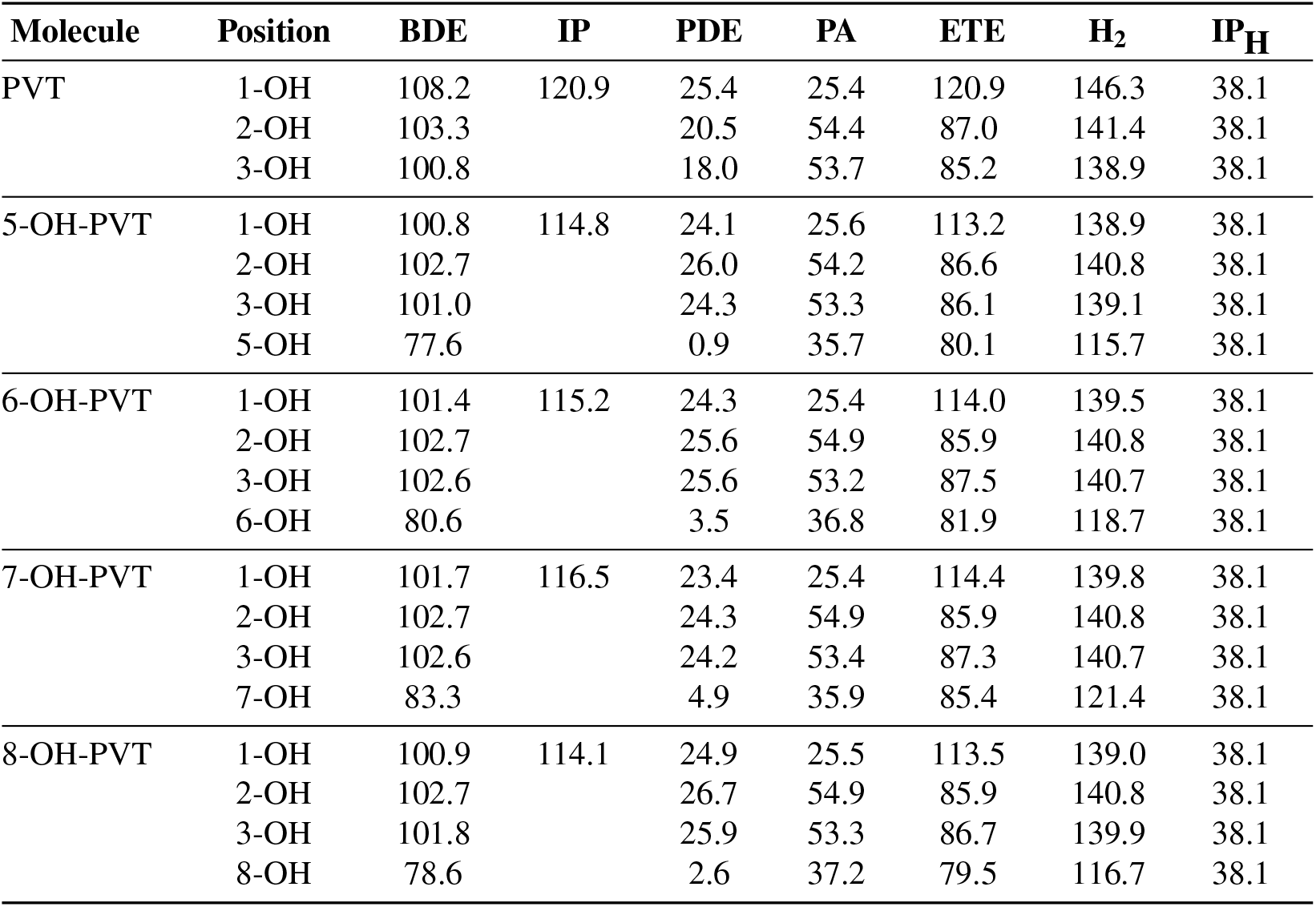
The enthalpies of reaction (in kcal/mol) needed to quantify the antioxidant activity of pitavastatin (PVT) and its hydroxylated derivatives in methanolic phase computed at *T* = 28.15 K. Notice that “combined” enthalpies IP + PDE and PA + ETE of the two-step mechanisms for a given molecular species are equal (*H*_2_ = IP + PDE = PA + ETE), and the difference *H*_2_ *−* BDE has the same value for all molecular species in a given environment (methanolic phase in the present cases), which coincides with the ionization enthalpy of the H-atom IPH (*H*_2_ *−* BDE =IPH) in given environment considered (methanolic phase in the present cases), obeying thereby the two theorems for antioxidation recently demonstrated [55].

As emphasized recently [53], it is impossible to unambiguously assessing the dominant mechanism of antioxidation merely comparing among themselves the various enthalpies of reaction characterizing the antioxidant (PVT in our case) without information for the free radical to be scavenged, e.g., its electron withdrawing power as well as on the reaction kinetics.

In the same vein, we should reiterate that the comparison between the combined enthalpies of reactions (BDE, IP + PDE, and PA + ETE) of the various mechanisms (HAT, SETPT, and SPLET, respectively) is meaningless [55]: irrespective of the specific antioxidant envisaged, the equality IP + PDE = PA + ETE(= *H*_2_) holds in general, and the value *H*_2_ common for the two two-step mechanism (SETET and SPLET) is always larger than the enthalpy of reaction BDE characterizing the single-step mechanism (HAT). Irrespective of the specific antioxidant and the molecular position of H atom abstraction, the difference *H*_2_ *−* BDE *>* 0 is equal to hydrogen’s ionization potential (IPH) in the medium (solvent) considered (*H*_2_ *−* BDE = IPH).

With this grain of salt, we can still remark that all the three OH positions possess bond dissociation enthalpies smaller than PVT’s ionization potential (BDE *<* IP). This makes the SETPT mechanism unlikely. Out of the three OH positions, position 1 (OH group of the carboxylic acid) possesses the largest bond dissociation enthalpy (BDE = 108.2 kcal/mol) but the smallest proton affinity (PA = 25.4 kcal/mol). Notwithstanding this small PA-value— which is not much larger that that computed for the lowest value of ascorbic acid (namely PA = 20.5 kcal/mol [54], a value computed at the present B3LYP/6-31+G(d,p)/IEFPCM level of theory)—, H-atom donation via an SPLET mechanism from position 1-OH is problematic; the enthalpy of the SPLET second step, which acts as a bottleneck, is higher (BDE = 108.2 kcal/mol *<* ETE = 120.9 kcal/mol).

### 3.3. Impact on Antioxidant Activity of a H Atom Replacement by a OH Group at Various Positions of the Quinoline Core

It is amply documented that chemical substituents significantly impact on the antiradical activity [86,93]. In vein with this general behavior, and in the light of the recent findings for the related atorvastatin molecule mentioned in Introduction [53], we systematically investigated the impact of replacing a hydrogen atom by a hydroxy group located in various positions on PVT’s aromatic rings. Results will be presented separately for replacements located in the heterocyclic quinoline and for replacements on the fluorophenyl ring.

In this subsection we will consider the effect of hydroxylation at positions 5, 6, 7, and 8 on the quinoline moiety. The labels employed are in accord to the standard atom numbering utilized for quinoline (Figure 1). The optimized geometries of all these (let us call them) Q-hydroxylated PVT derivatives as well as the pertaining HOMO and LUMO spatial distributions are depicted in Figures 2 to 5. For comparative purposes, their counterparts for the isolated quinoline molecules with H atoms replaced by OH groups are also presented in these figures.

**Figure 2.**
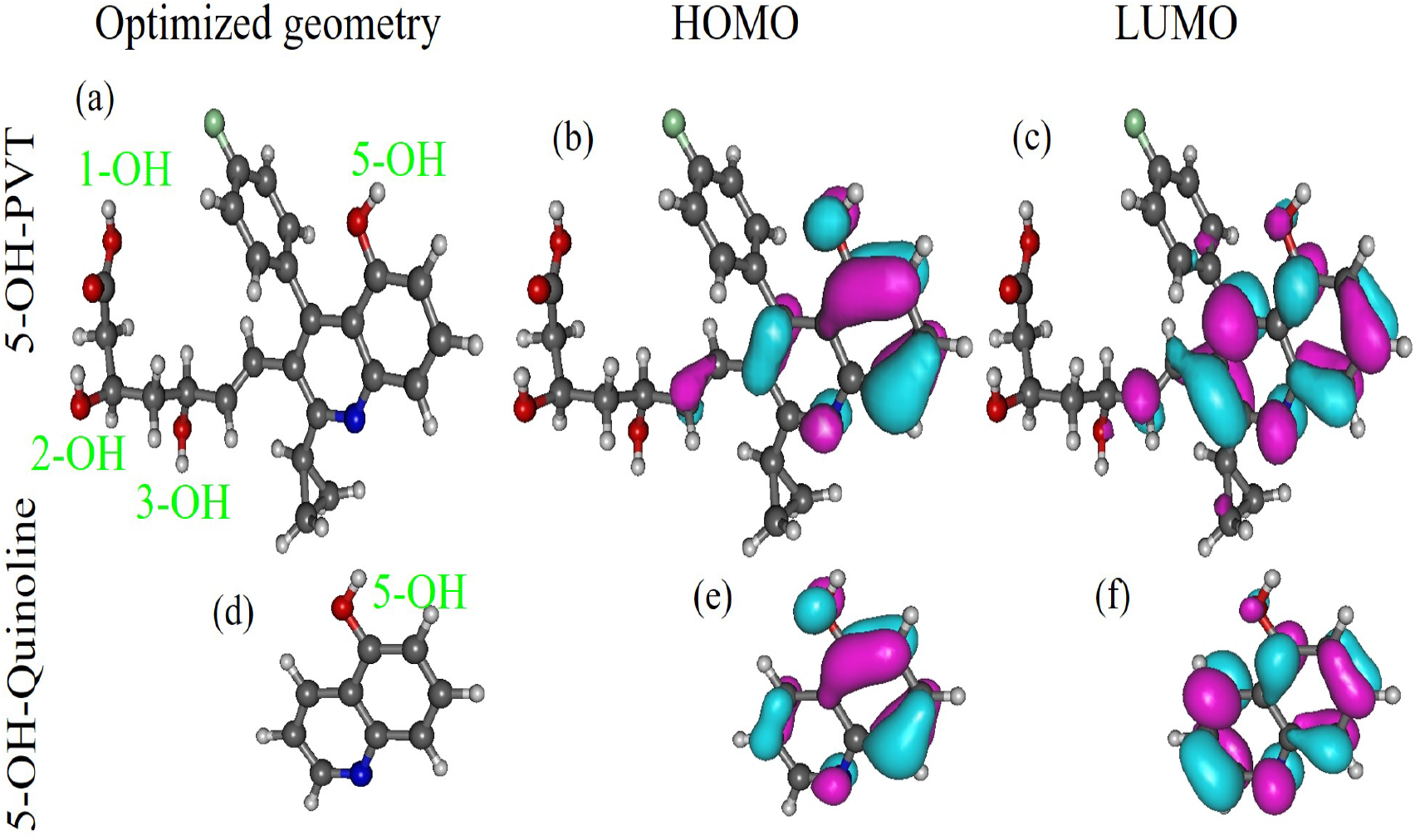
(a) The optimized geometry of the PVT molecule wherein the H atom at position 5 on the quinoline core is replaced by an OH group and the corresponding (a) HOMO and (b) LUMO spatial distributions. Their counterparts for the isolated quinoline molecule are depicted in (c), (d), and (e), respectively. Figures generated using Gabedit 2.5.1 [92].

**Figure 3.**
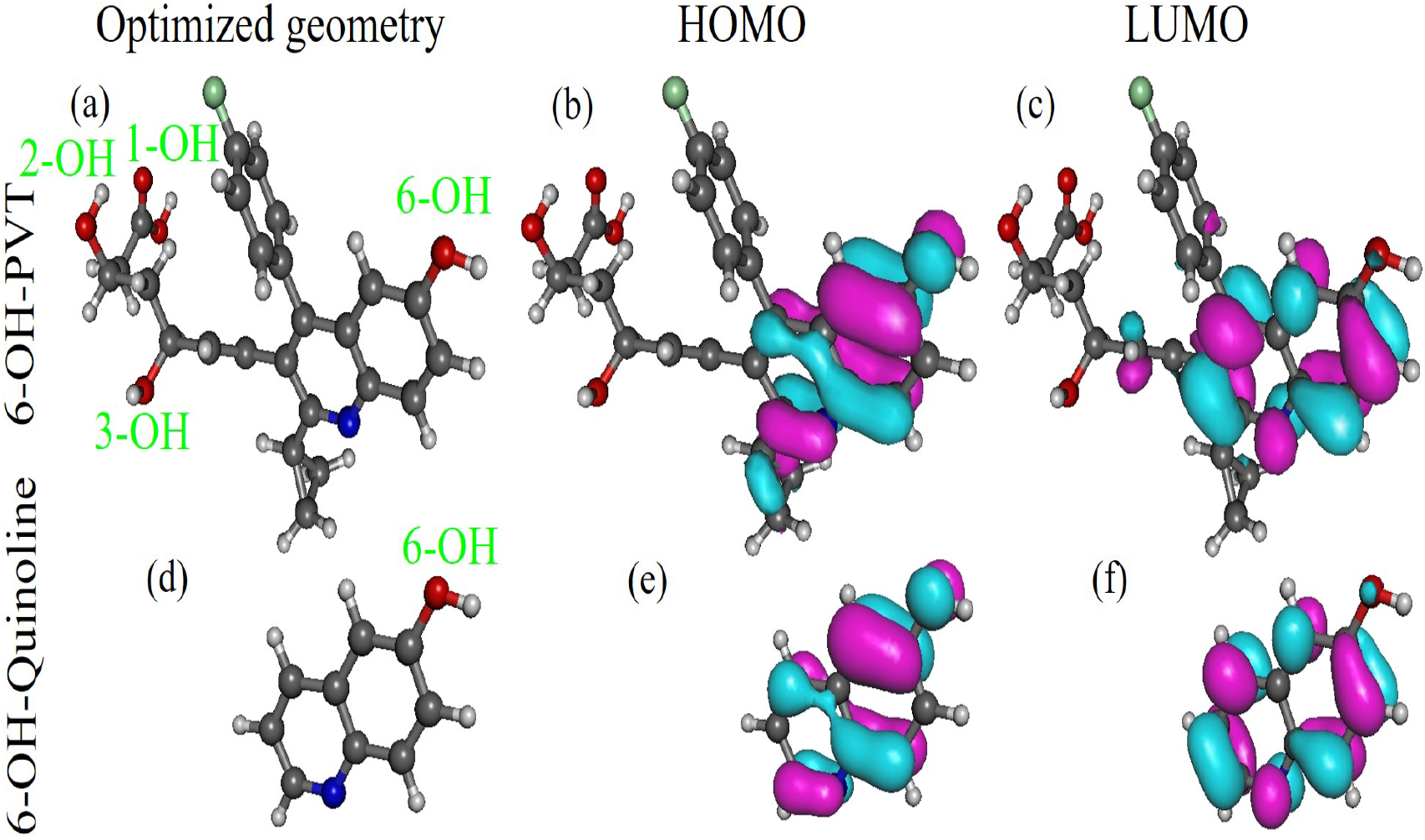
(a) The optimized geometry of the PVT molecule wherein the H atom at position 6 on the quinoline core is replaced by an OH group and the corresponding (a) HOMO and (b) LUMO spatial distributions. Their counterparts for the isolated quinoline molecule are depicted in (c), (d), and (e), respectively. Figures generated using Gabedit 2.5.1 [92].

**Figure 4.**
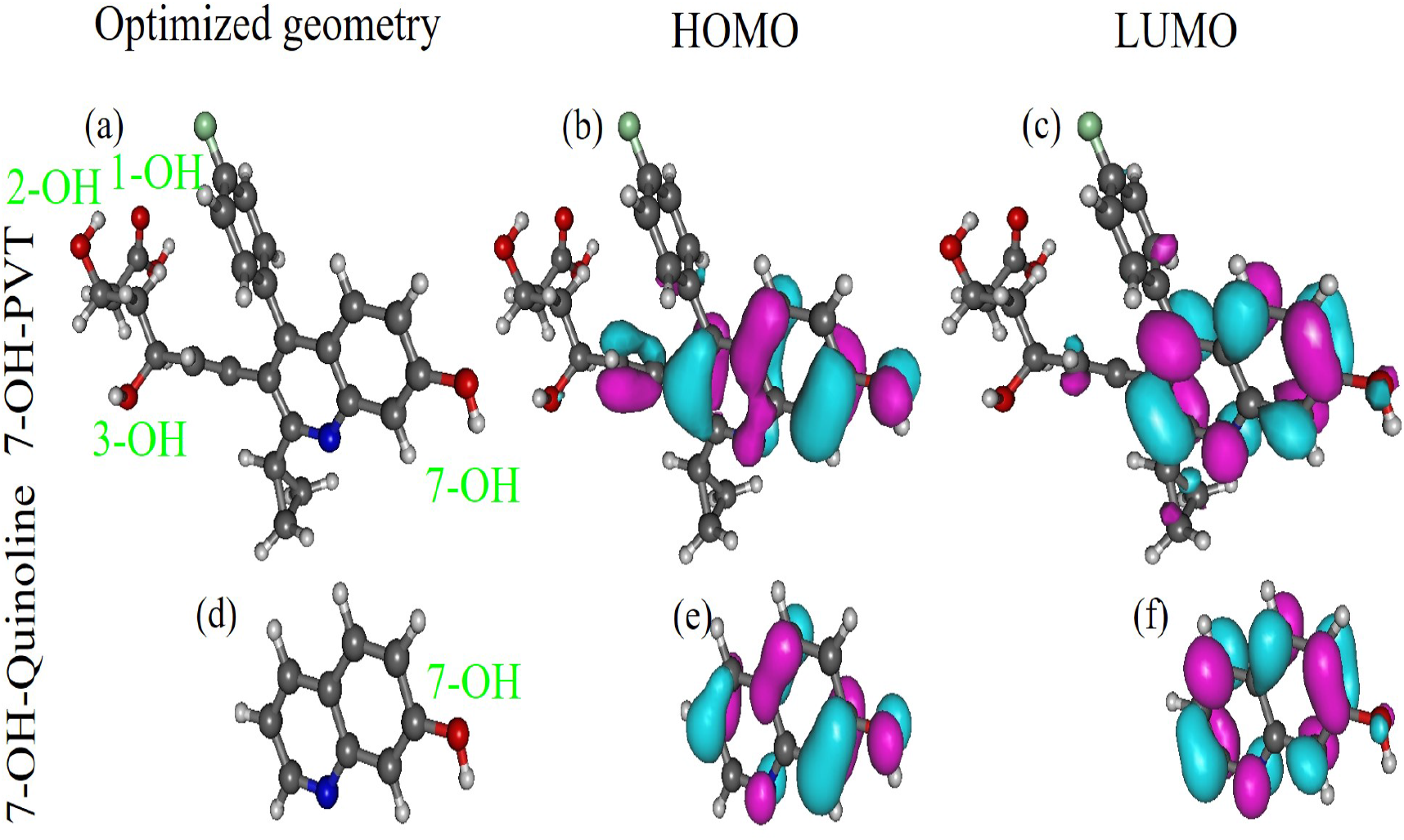
(a) The optimized geometry of the PVT molecule wherein the H atom at position 7 on the quinoline core is replaced by an OH group and the corresponding (a) HOMO and (b) LUMO spatial distributions. Their counterparts for the isolated quinoline molecule are depicted in (c), (d), and (e), respectively. Figures generated using Gabedit 2.5.1 [92].

**Figure 5.**
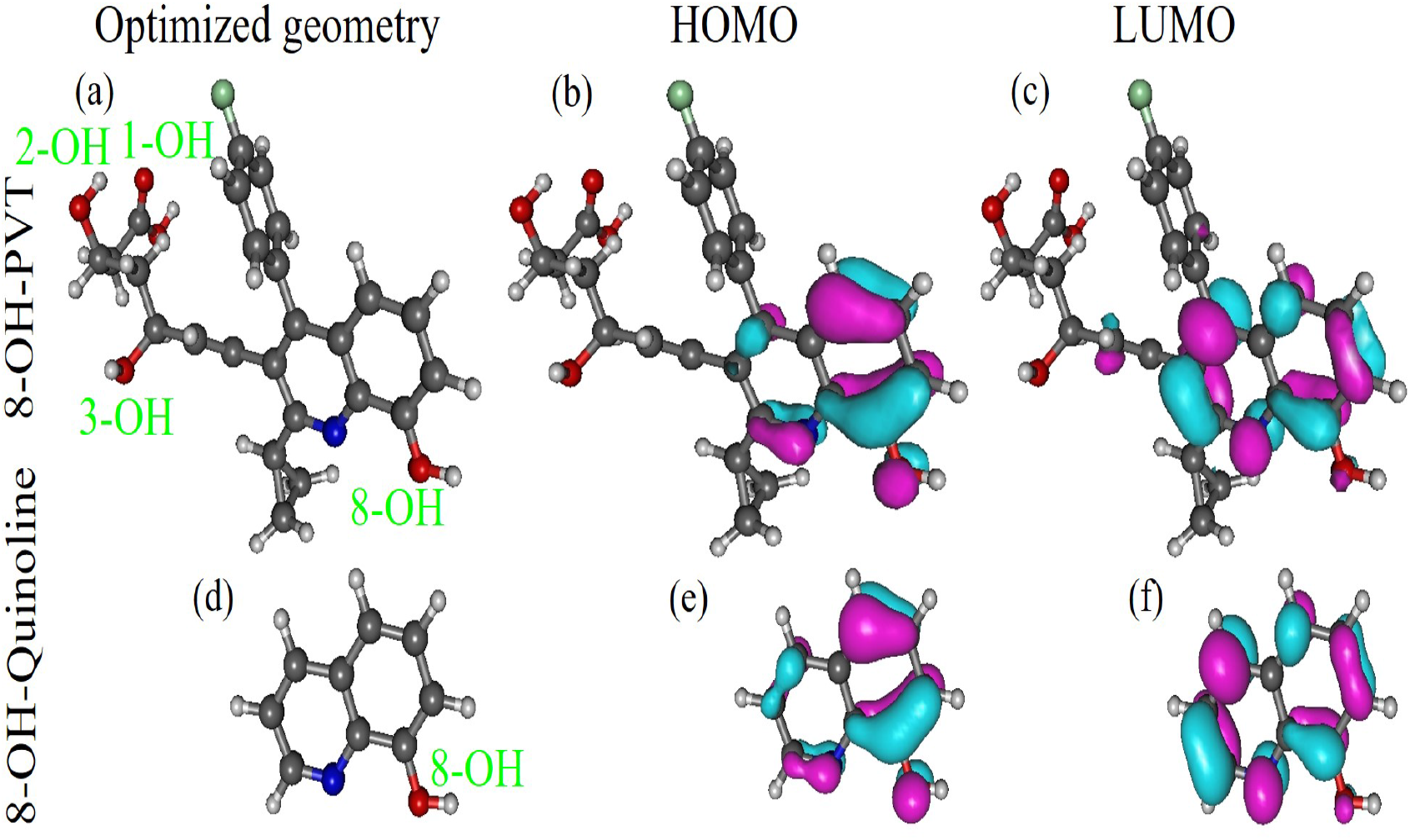
(a) The optimized geometry of the PVT molecule wherein the H atom at position 8 on the quinoline core is replaced by an OH group and the corresponding (a) HOMO and (b) LUMO spatial distributions. Their counterparts for the isolated quinoline molecule are depicted in (c), (d), and (e), respectively. Figures generated using Gabedit 2.5.1 [92].

Our results for the enthalpies of reaction quantifying the antioxidant properties at all these four positions (5, 6, 7, 8) available for hydrogen atom abstraction in PVT’s hydroxylated metabolites are presented in Table 1 along with those for the parent PVT molecule. Most importantly, whether at position 5, 6, 7, or 8, the extra fourth OH group attached to the quinoline moiety possesses a bond dissociation energy BDE substantially smaller than those of PVT (at positions 1, 2, and 3, BDE *>* 100 kcal/mol). Out of these Q-hydroxylated metabolites, 5-OH-PVT possesses the smallest BDE value (BDE = 77.6, kcal/mol), followed by 8-OH-PVT (BDE = 78.6, kcal/mol).

For all Q-hydroxylated derivatives, the extra OH group attached to the quinoline core has a BDE value smaller than the next lowest BDE value by amounts ranging from 18.4 kcal/mol to 23.2 kcal/mol. Interestingly, this next lowest BDE is in all cases associated with the 1-OH group of the carboxylic acid, in contrast to the parent PVT wherein the group 1-OH possesses the largest BDE. The overall reduction in BDE at all positions (1, 2, and 3) characterizing the parent PVT molecules represents another significant effect of hydroxylating the quinoline core.

The impact of hydroxylation on the ionization potential IP is similar to that on the BDE values at the positions 1, 2, and 3 inherited from PVT. All Q-hydroxylated derivatives have values of IP smaller than the value for the parent PVT; the computed differences— smaller than those for BDE’s by a factor of 3 *−* 4— range from 4.4 kcal/mol to 6.8 kcal/mol.

The aforementioned behavior contrasts to the effect of the additional OH group on the heterolytic O-H bond cleavage. Proton affinities (PA) at positions 1, 2, and 3 are virtually unaffected. Being smaller than 1 kcal/mol (“chemical accuracy”), the computed differences in the corresponding values of PA are irrelevant. As for the positions on the quinoline core, all PA values are larger by about 10 kcal/mol than the lowest PA value, namely the value at position 1 (carboxylic acid). If heterolytic O-H bond cleavage were to occur in Q-hydroxylated PVT derivatives, it would rather occur within the carboxylic acid group. The bottom line is that, if SPLET were the dominant mechanism, hdroxylated PVT metabolites would not be better free radical scavengers than the parent PVT.

Comparison between the impact on the antioxidant activity of hydroxy substitutions at various positions on PVT’s quinoline moiety with those on the isolated quinoline molecule is also interesting. Numerical values for the antioxidant properties of the hydroxylated derivatives of the isolated quinoline molecule along with those characterizing the pertaining positions on the quinoline moiety from the Q-hydroxylated PVT metabolites are presented in Table 2. To facilitate this comparison, we additionally depict the corresponding values of BDE, IP, PDE, PA, and ETE in Figures 6 to 10. Inspection of Figures 6b to 10b reveals that the values of BDE, IP, and ETE of the hydroxy groups in isolated quinoline are somewhat larger than those of the Q-hydroxylated PVT metabolites, while the opposite holds true for the values of PDE and PA. Despite these rather minor differences, the antioxidant properties of the Q-hydroxylated PVT metabolites and those of the hydroxylated quinoline species are well correlated. The good correlation between the values of BDE, IP, PA, and ETE (expressed by the large values of *R*^2^) is clearly seen in Figures 6c, 7c, 9c, and 10c, respectively. The significantly poorer correlation for PDE’s depicted in Figure 8c should be related to the small values of PDE (much smaller than the other entahlpies of reaction), for which the aforementioned “chemical accuracy” cannot be expected.

**Table 2.** The enthalpies of reaction (in kcal/mol) needed to quantify the antioxidant activity of quinoline’s hydroxylated derivatives in methanolic phase computed at *T* = 28.15 K. Notice that “combined” enthalpies IP+PDE and PA+ETE of the two-step mechanisms for a given molecular species are equal (*H*_2_ = IP+PDE = PA + ETE), and the difference *H*_2_ *−* BDE has the same value for all molecular species in a given environment (methanolic phase in the present cases), which coincides with the ionization enthalpy of the H-atom IPH (*H*_2_ *−* BDE =IPH) in given environment considered (methanolic phase in the present cases), obeying thereby the two theorems for antioxidation recently demonstrated [55]

| Molecule | BDE | IP | PDE | PA | ETE | $H_2$ | $IP_H$ |
| --- | --- | --- | --- | --- | --- | --- | --- |
| 5-OH-PVT | 77.6 | 114.8 | 0.9 | 35.7 | 80.1 | 115.7 | 38.1 |
| 5-OH-Q | 79.2 | 119.7 | -2.5 | 34.5 | 82.7 | 117.3 | 38.1 |
| 6-OH-PVT | 80.6 | 115.2 | 3.5 | 36.8 | 81.9 | 118.7 | 38.1 |
| 6-OH-Q | 82.6 | 122.0 | -1.3 | 35.7 | 85.0 | 120.7 | 38.1 |
| 7-OH-PVT | 83.3 | 116.5 | 4.9 | 35.9 | 85.4 | 121.4 | 38.1 |
| 7-OH-Q | 84.2 | 124.0 | -1.7 | 35.6 | 86.7 | 122.2 | 38.1 |
| 8-OH-PVT | 78.6 | 114.1 | 2.6 | 37.2 | 79.5 | 116.7 | 38.1 |
| 8-OH-Q | 79.7 | 119.1 | -1.3 | 36.3 | 81.4 | 117.8 | 38.1 |

**Figure 6.**
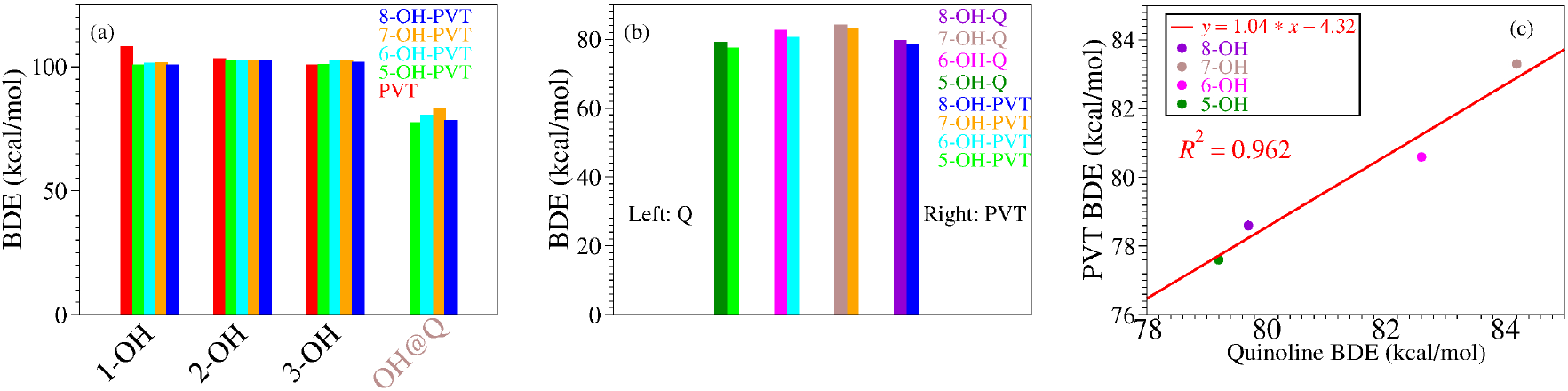
Panel a presents bond dissociation enthalpies BDE computed for the PVT molecule and its Q-hydroxylated derivatives. Positions 1, 2, and 3 denote the positions of the OH group in the parent PVT, nOH@Q (*n* = 5, 6, 7, 8) refers to the additional OH group replacing an H atom on the quinoline moiety of the Q-hydroxylated PVT species. In panels b and c, the nOH@Q-values (rightmost bars of the histogram in panel a) are depicted along the corresponding bond dissociation enthalpies of the isolated quinoline (Q) molecule wherein the H atom at positions 5, 6, 7, and 8 were replaced by an OH group. The large value of *R*^2^ in panel c visualizes the good correlation between the BDE values for the various hydroxylated positions on the PVT’s quinoline core and the BDE values corresponding to the isolated quinoline molecule.

**Figure 7.**
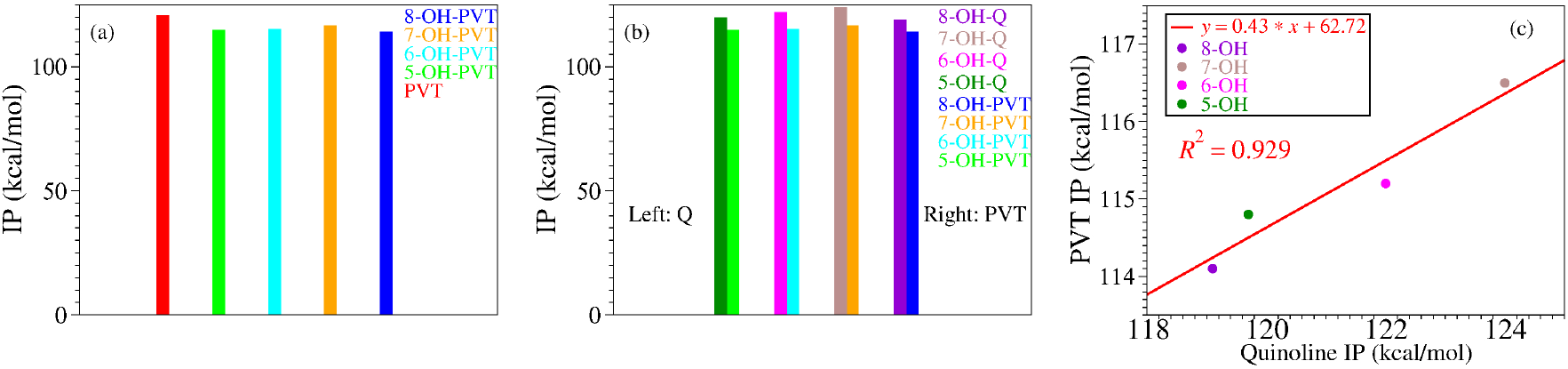
Ionization potentials IP computed for the PVT molecule and the various Q-hydroxylated PVT derivatives (panel a). In panels b and c, the latter are depicted along the corresponding ionization potentials of the isolated quinoline (Q) molecule wherein the H atom at positions 5, 6, 7, and 8 were replaced by an OH group. The large value of *R*^2^ in panel c visualizes the good correlation between the IP values for the various hydroxylated positions on the PVT’s quinoline core and the IP values corresponding to the isolated quinoline molecule.

**Figure 8.**
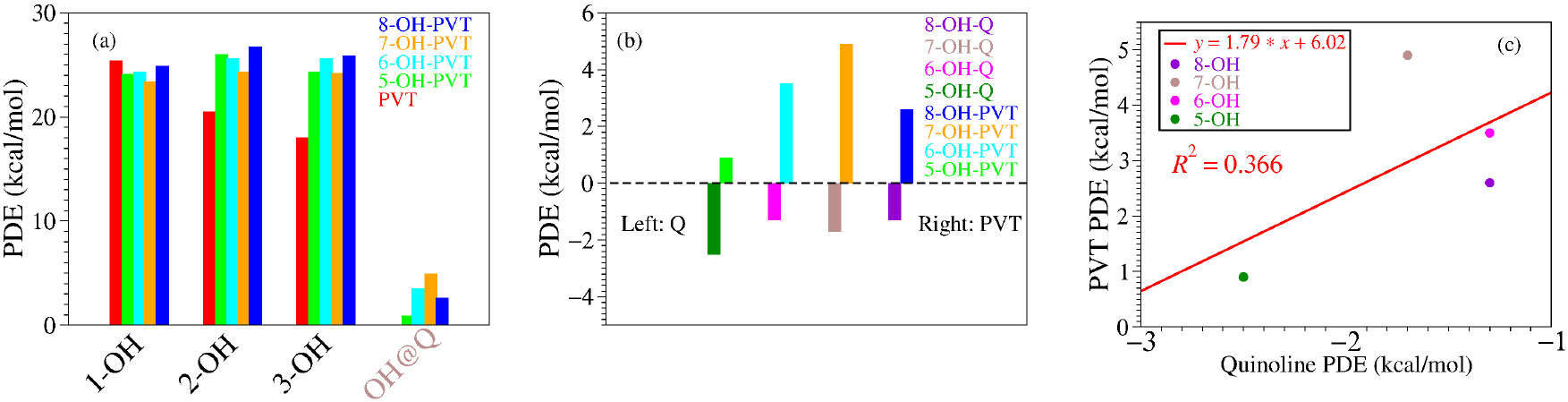
Panel a presents proton dissociation enthalpies PDE computed for the PVT molecule and its Q-hydroxylated derivatives. Positions 1, 2, and 3 denote the positions of the OH group in the parent PVT, nOH@Q (*n* = 5, 6, 7, 8) refers to the additional OH group replacing an H atom on the quinoline moiety of the Q-hydroxylated PVT species. In panels b and c, the nOH@Q-values (rightmost bars of the histogram in panel a) are depicted along the corresponding bond dissociation enthalpies of the isolated quinoline (Q) molecule wherein the H atom at positions 5, 6, 7, and 8 were replaced by an OH group. As elaborated in the main text, the value of *R*^2^ in panel c— much smaller than for the other enthalpies of reaction shown in Figures 6, 7, 9, and 10— does not necessarily express a poorer correlation between the PDE values for the various hydroxylated positions on the PVT’s quinoline core and the PDE values corresponding to the isolated quinoline molecule.

**Figure 9.**
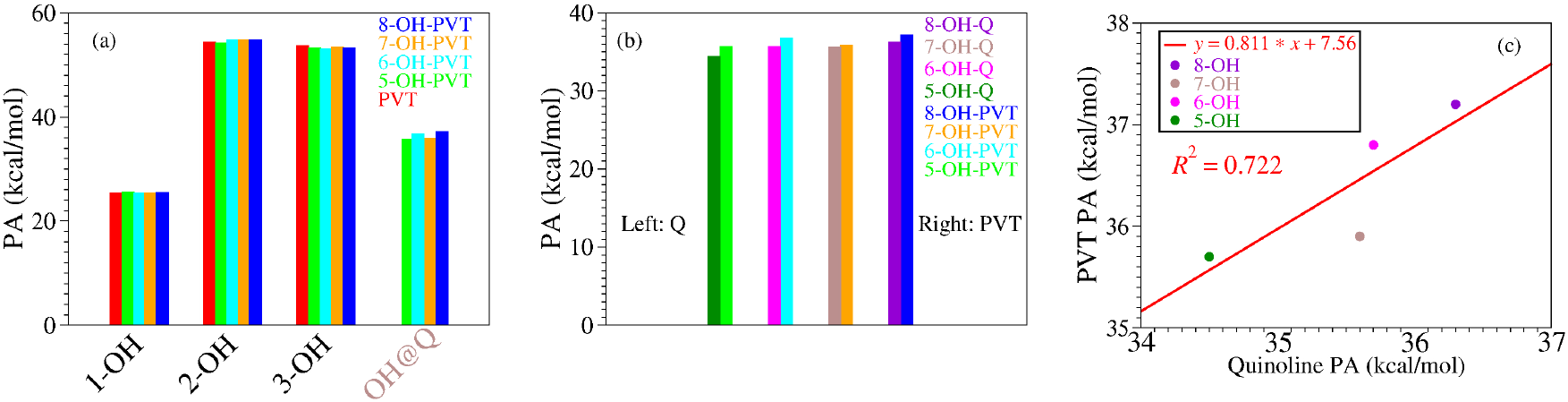
Panel a presents proton affinities PA computed for the PVT molecule and its Q-hydroxylated derivatives. Positions 1, 2, and 3 denote the positions of the OH group in the parent PVT, nOH@Q (*n* = 5, 6, 7, 8) refers to the additional OH group replacing an H atom on the quinoline moiety of the Q-hydroxylated PVT species. In panels b and c, the nOH@Q-values (rightmost bars of the histogram in panel a) are depicted along the corresponding proton affinities of the isolated quinoline (Q) molecule wherein the H atom at positions 5, 6, 7, and 8 were replaced by an OH group. The value of *R*^2^ in panel c visualizes a reasonable correlation between the PA values for the various hydroxylated positions on the PVT’s quinoline core and the PA values corresponding to the isolated quinoline molecule.

**Figure 10.**
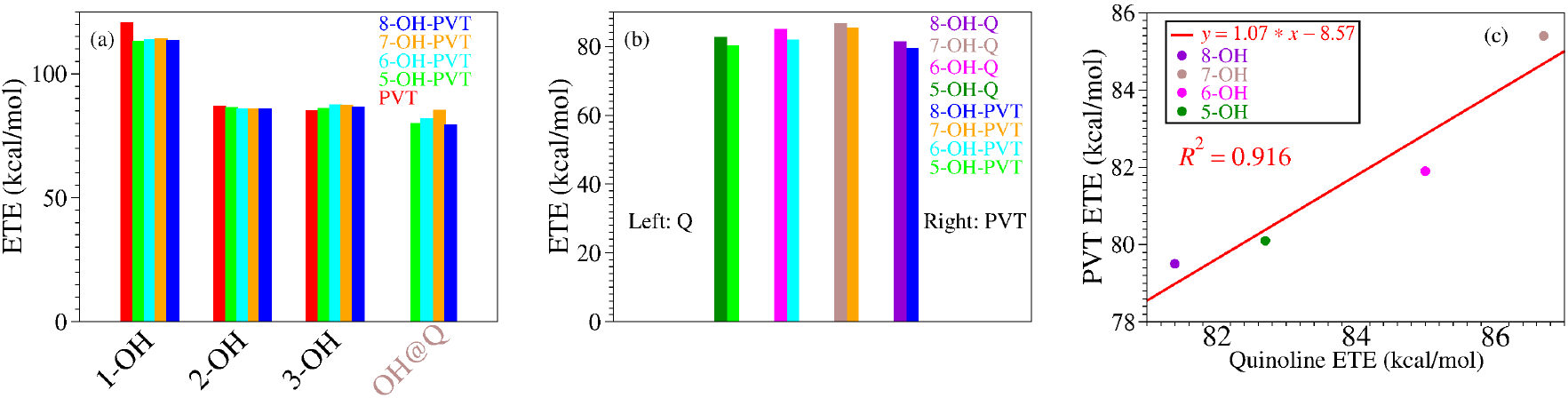
Panel a presents electron transfer enthalpies ETE computed for the PVT molecule and its Q-hydroxylated derivatives. Positions 1, 2, and 3 denote the positions of the OH group in the parent PVT, nOH@Q (*n* = 5, 6, 7, 8) refers to the additional OH group replacing an H atom on the quinoline moiety of the Q-hydroxylated PVT species. In panels b and c, the nOH@Q-values (rightmost bars of the histogram in panel a) are depicted along the corresponding electron transfer enthalpies of the isolated quinoline (Q) molecule wherein the H atom at positions 5, 6, 7, and 8 were replaced by an OH group. The large value of *R*^2^ in panel c visualizes the good correlation between the ETE values for the various hydroxylated positions on the PVT’s quinoline core and the ETE values corresponding to the isolated quinoline molecule.

### 3.4. Impact on Antioxidant Activity of a H Atom Replacement by a OH Group at Various Positions of the Fluorophenyl Ring

The hydroxylated derivatives resulting by replacing a hydrogen atom attached to the fluorophenyl ring (let us call them F-hydroxylated PVT derivatives) at their optimum geometry and their FMO distributions are depicted in Figures 11 to 14. For comparison purposes, the fluorinated phenol species corresponding to them are also shown there.

**Figure 11.**
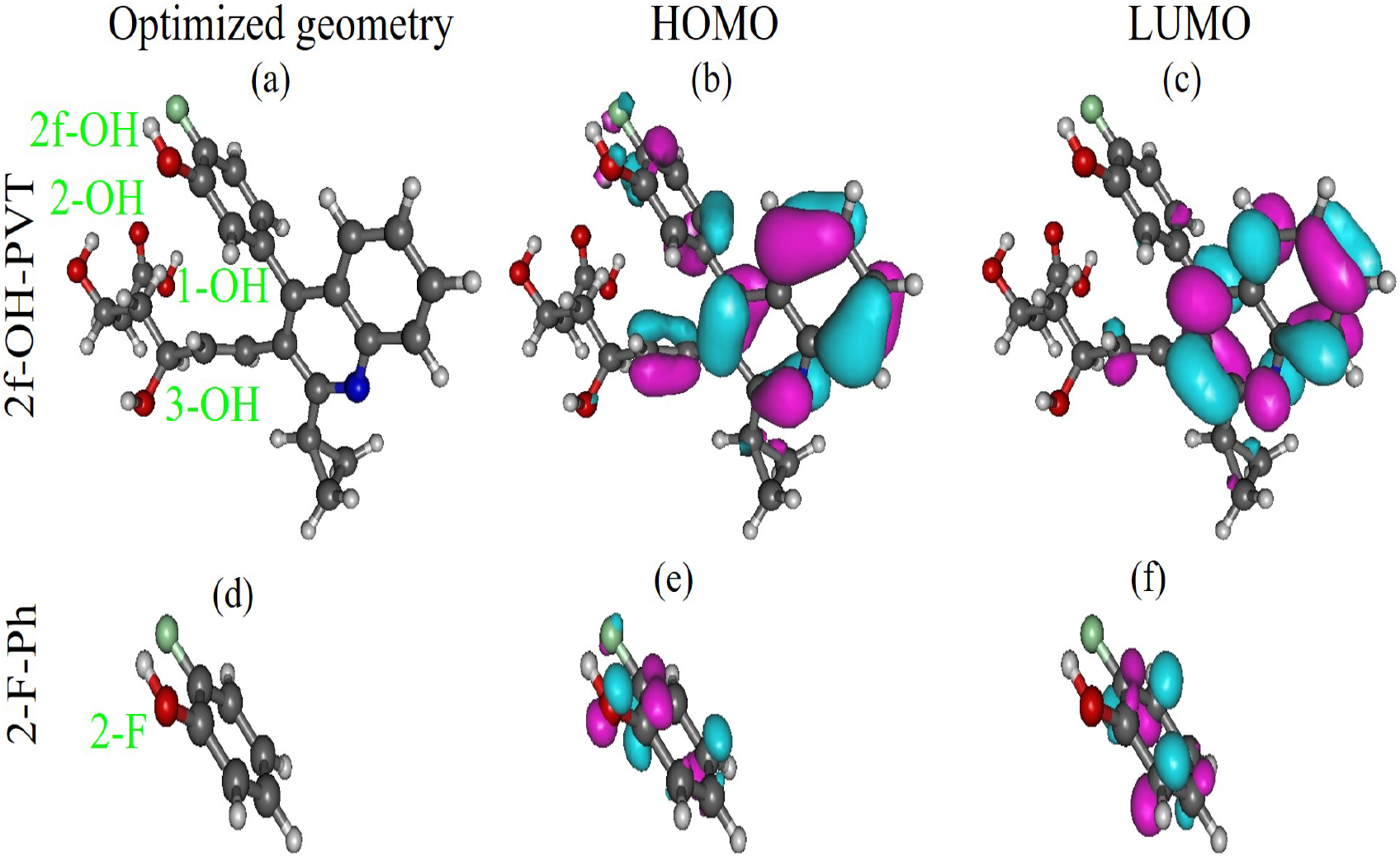
(a) The optimized geometry of the PVT molecule wherein the H atom at position 2f on the fluorophenyl moiety is replaced by an OH group and the corresponding (a) HOMO and (b) LUMO spatial distributions. Their counterparts for the isolated 2-F fluorophenol molecule are depicted in (c), (d), and (e), respectively. Figures generated using Gabedit 2.5.1 [92].

**Figure 12.**
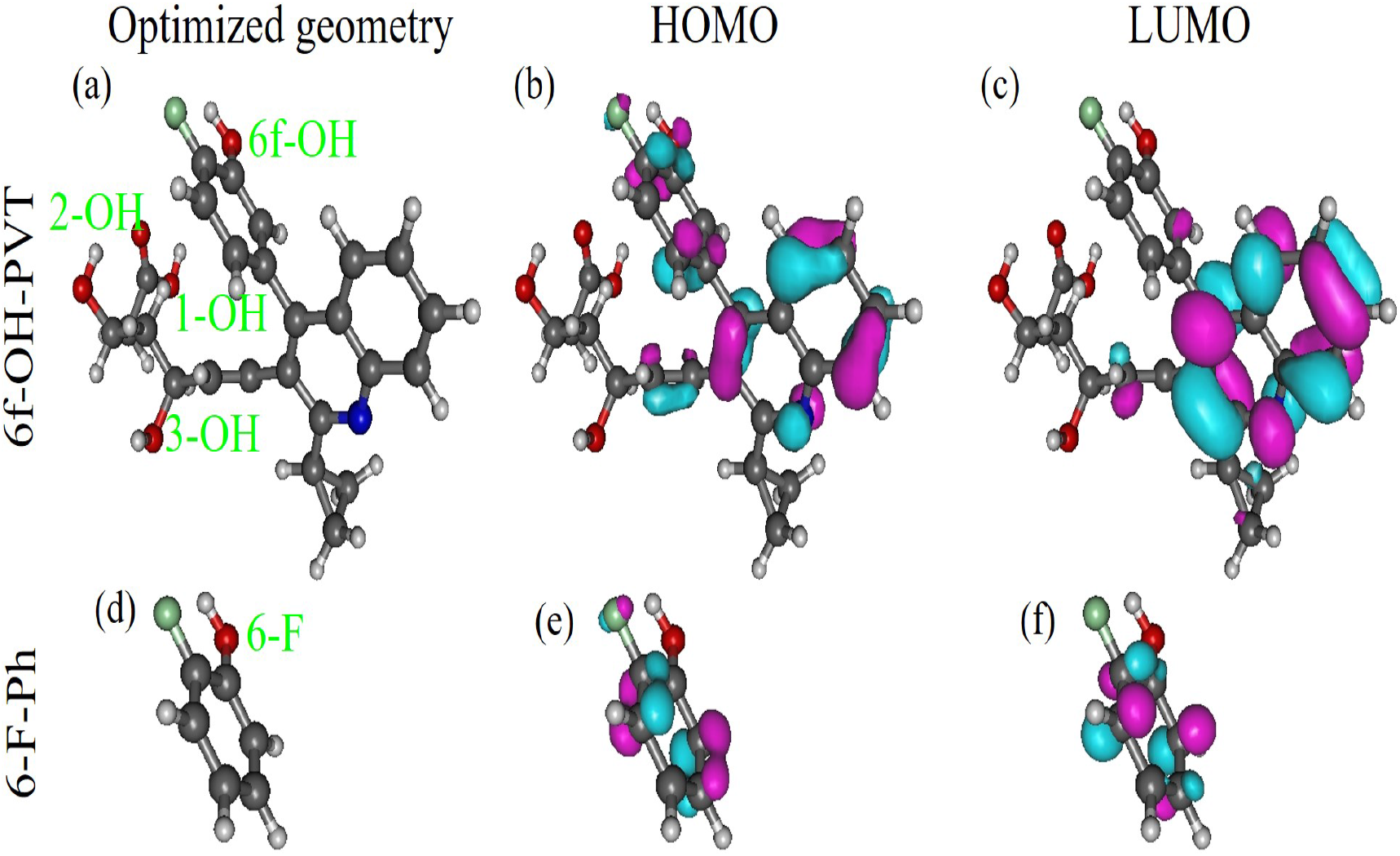
(a) The optimized geometry of the PVT molecule wherein the H atom at position 6f on the fluorophenyl moiety is replaced by an OH group and the corresponding (a) HOMO and (b) LUMO spatial distributions. Their counterparts for the isolated 6-F (identical to 2-F) fluorophenol molecule are depicted in (c), (d), and (e), respectively. Figures generated using Gabedit 2.5.1 [92].

**Figure 13.**
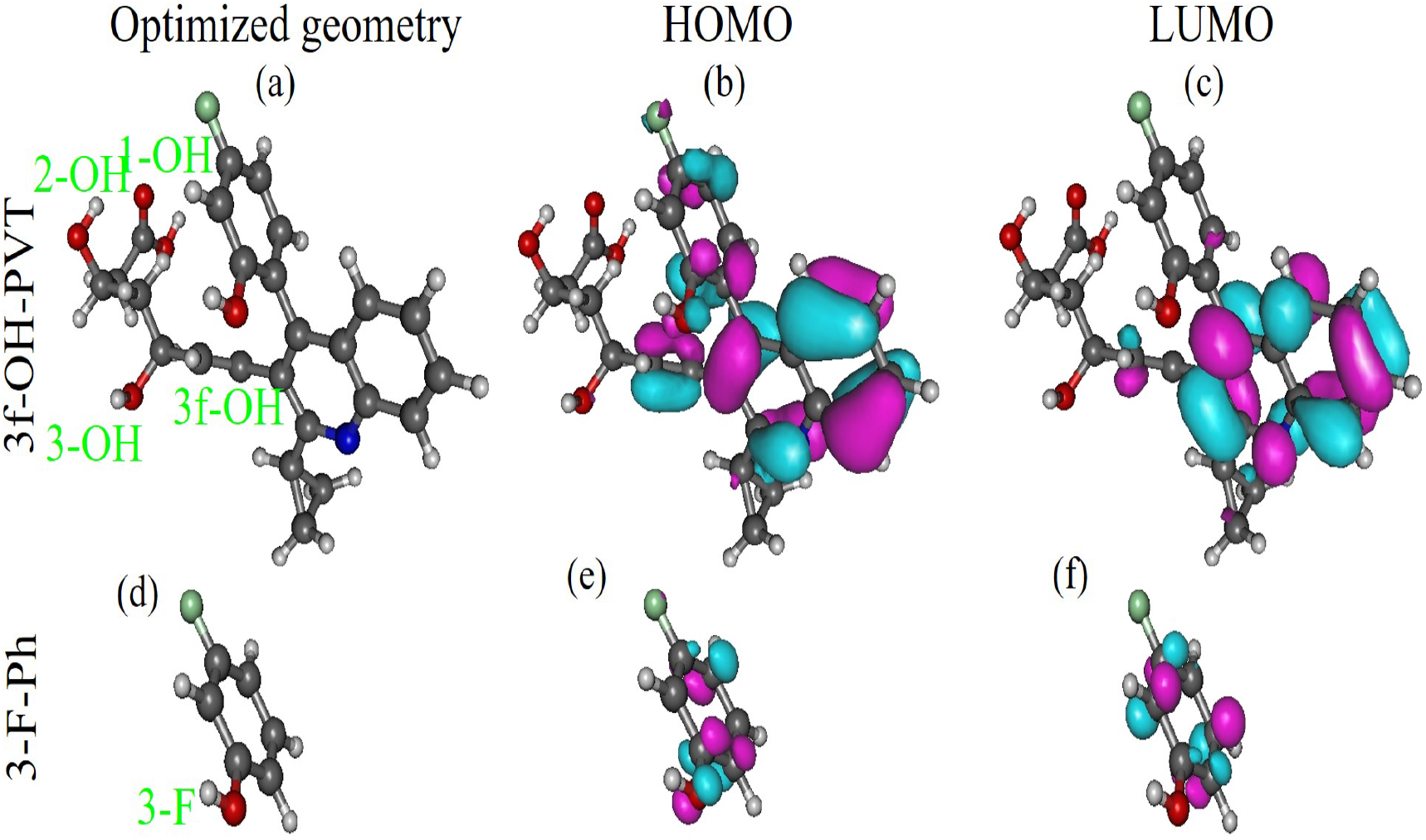
(a) The optimized geometry of the PVT molecule wherein the H atom at position 3f on the fluorophenyl moiety is replaced by an OH group and the corresponding (a) HOMO and (b) LUMO spatial distributions. Their counterparts for the isolated 3-F fluorophenol molecule are depicted in (c), (d), and (e), respectively. Figures generated using Gabedit 2.5.1 [92].

**Figure 14.**
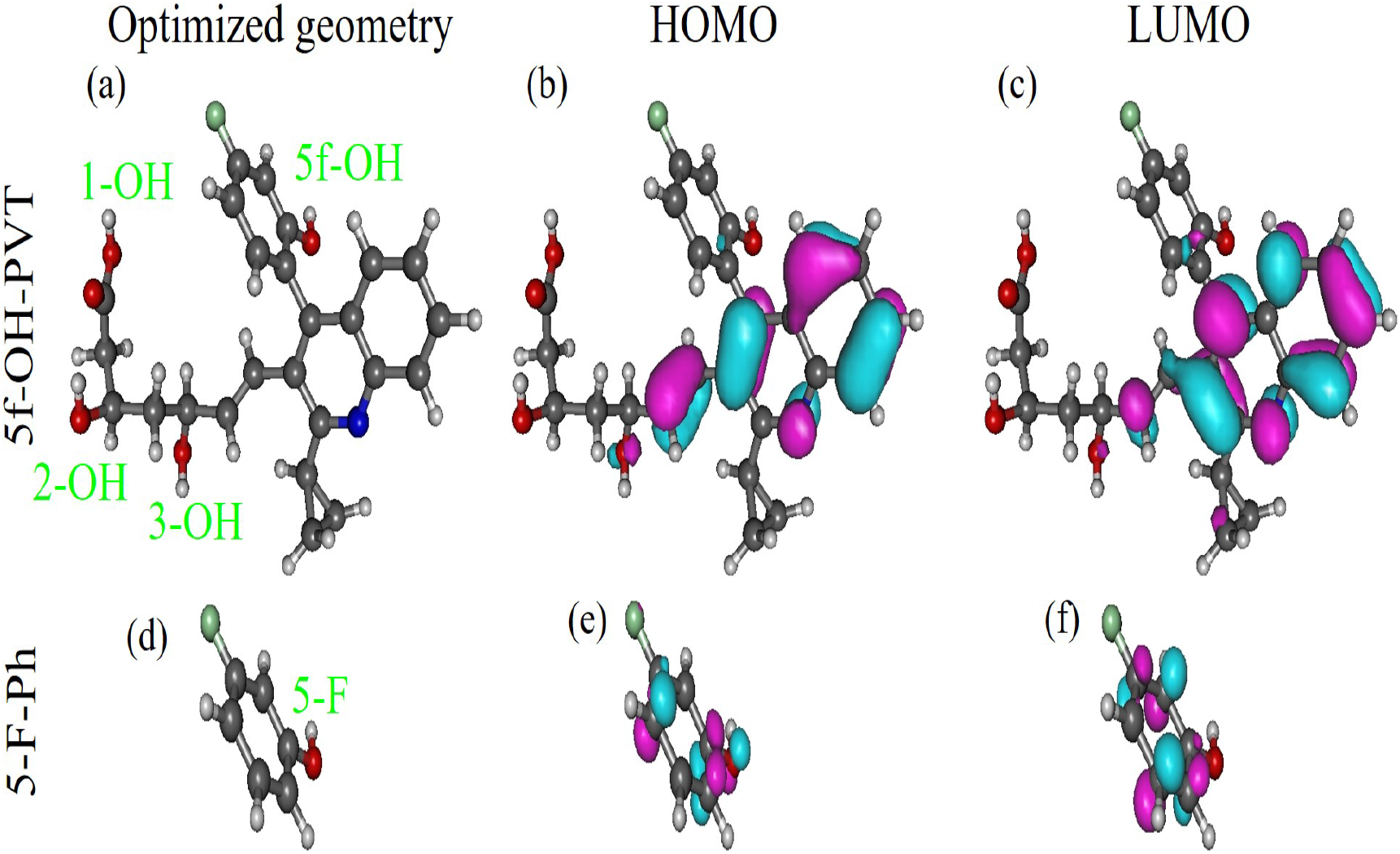
(a) The optimized geometry of the PVT molecule wherein the H atom at position 5f on the fluorophenyl moiety is replaced by an OH group and the corresponding (a) HOMO and (b) LUMO spatial distributions. Their counterparts for the isolated 5-F (identical to 3-F) fluorophenol molecule are depicted in (c), (d), and (e), respectively. Figures generated using Gabedit 2.5.1 [92].

In accord with chemical intuition, calculations confirm that neither the FMO distributions nor the enthalpies of reaction pertaining to positions 1, 2, and 3 of the parent PVT molecule are notably affected by the OH-group replacing an hydrogen atom of the fluorophenyl ring; see Table 3.

**Table 3.** The enthalpies of reaction (in kcal/mol) needed to quantify the antioxidant activity of F-hydroxylated PVT’s derivatives at position 1 (corresponding to the carboxylic group) in methanolic phase computed at *T* = 28.15 K. Notice that “combined” enthalpies IP + PDE and PA + ETE of the two-step mechanisms for a given molecular species are equal (*H*_2_ = IP + PDE = PA + ETE), and the difference *H*_2_ *−* BDE has the same value for all molecular species in a given environment (methanolic phase in the present cases), which coincides with the ionization enthalpy of the H-atom IPH (*H*_2_ *−* BDE =IPH) in given environment considered (methanolic phase in the present cases), obeying thereby the two theorems for antioxidation recently demonstrated [55].

| Molecule | Position | BDE | IP | PDE | PA | ETE | $H_2$ | $IP_H$ |
| --- | --- | --- | --- | --- | --- | --- | --- | --- |
| PVT | 1-OH | 108.2 | 120.9 | 25.4 | 25.4 | 120.9 | 146.3 | 38.1 |
| 6f-OH-PVT | 1-OH | 108.2 | 121.0 | 25.3 | 25.5 | 120.8 | 146.3 | 38.1 |
| 5f-OH-PVT | 1-OH | 108.4 | 121.5 | 25.1 | 25.6 | 120.9 | 146.5 | 38.1 |
| 5-OH-PVT | 1-OH | 100.8 | 114.8 | 24.1 | 25.6 | 113.2 | 138.9 | 38.1 |
| 6-OH-PVT | 1-OH | 101.4 | 115.2 | 24.3 | 25.4 | 114.0 | 139.5 | 38.1 |
| 7-OH-PVT | 1-OH | 101.7 | 116.5 | 23.4 | 25.4 | 114.4 | 139.8 | 38.1 |
| 8-OH-PVT | 1-OH | 100.9 | 114.1 | 24.9 | 25.5 | 113.5 | 139.0 | 38.1 |

The relevant enthalpies of reaction of the corresponding F-hydroxylated PVT derivatives characterizing the antioxidant activity of the extra OH group attached to the fluorophenyl ring are collected in Table 4 and depicted in Figures 15. The fact that the F-hydroxylated PVT derivatives have values of the ionization potential considerably larger than those of the fluorinated phenol species while being closed to those of the parent PVT is directly related to the fact that, in contrast to the quinoline moiety (cf. Section 3.3), the fluorinated phenol moiety negligibly contributes to the HOMO of the F-hydoxylated PVT derivatives.

**Table 4.**
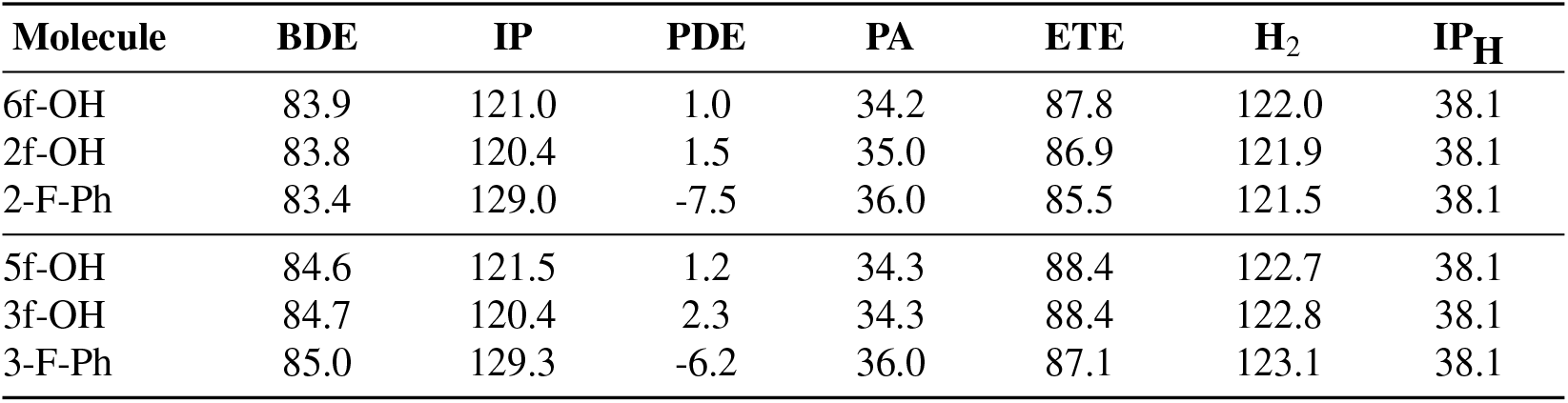
The enthalpies of reaction (in kcal/mol) needed to quantify the antioxidant activity of F-hydroxylated PVT’s derivatives computed in methanolic phase at *T* = 28.15 K for various positions (nf, *n* = 2, 3, 5, 6, labels as in Figures 12 to 14). Their counterparts for the isolated fluorophenol molecules (n-F-Ph, *n* = 2, 3) are also included. Notice that “combined” enthalpies IP + PDE and PA + ETE of the two-step mechanisms for a given molecular species are equal (*H*_2_ = IP + PDE = PA + ETE), and the difference *H*_2_ *−* BDE has the same value for all molecular species in a given environment (methanolic phase in the present cases), which coincides with the ionization enthalpy of the H-atom IPH (*H*_2_ *−* BDE =IPH) in given environment considered (methanolic phase in the present cases), obeying thereby the two theorems for antioxidation recently demonstrated [55].

| Molecule | BDE | IP | PDE | PA | ETE | $H_2$ | $IP_H$ |
| --- | --- | --- | --- | --- | --- | --- | --- |
| 6f-OH | 83.9 | 121.0 | 1.0 | 34.2 | 87.8 | 122.0 | 38.1 |
| 2f-OH | 83.8 | 120.4 | 1.5 | 35.0 | 86.9 | 121.9 | 38.1 |
| 2-F-Ph | 83.4 | 129.0 | -7.5 | 36.0 | 85.5 | 121.5 | 38.1 |
| 5f-OH | 84.6 | 121.5 | 1.2 | 34.3 | 88.4 | 122.7 | 38.1 |
| 3f-OH | 84.7 | 120.4 | 2.3 | 34.3 | 88.4 | 122.8 | 38.1 |
| 3-F-Ph | 85.0 | 129.3 | -6.2 | 36.0 | 87.1 | 123.1 | 38.1 |

**Figure 15.**
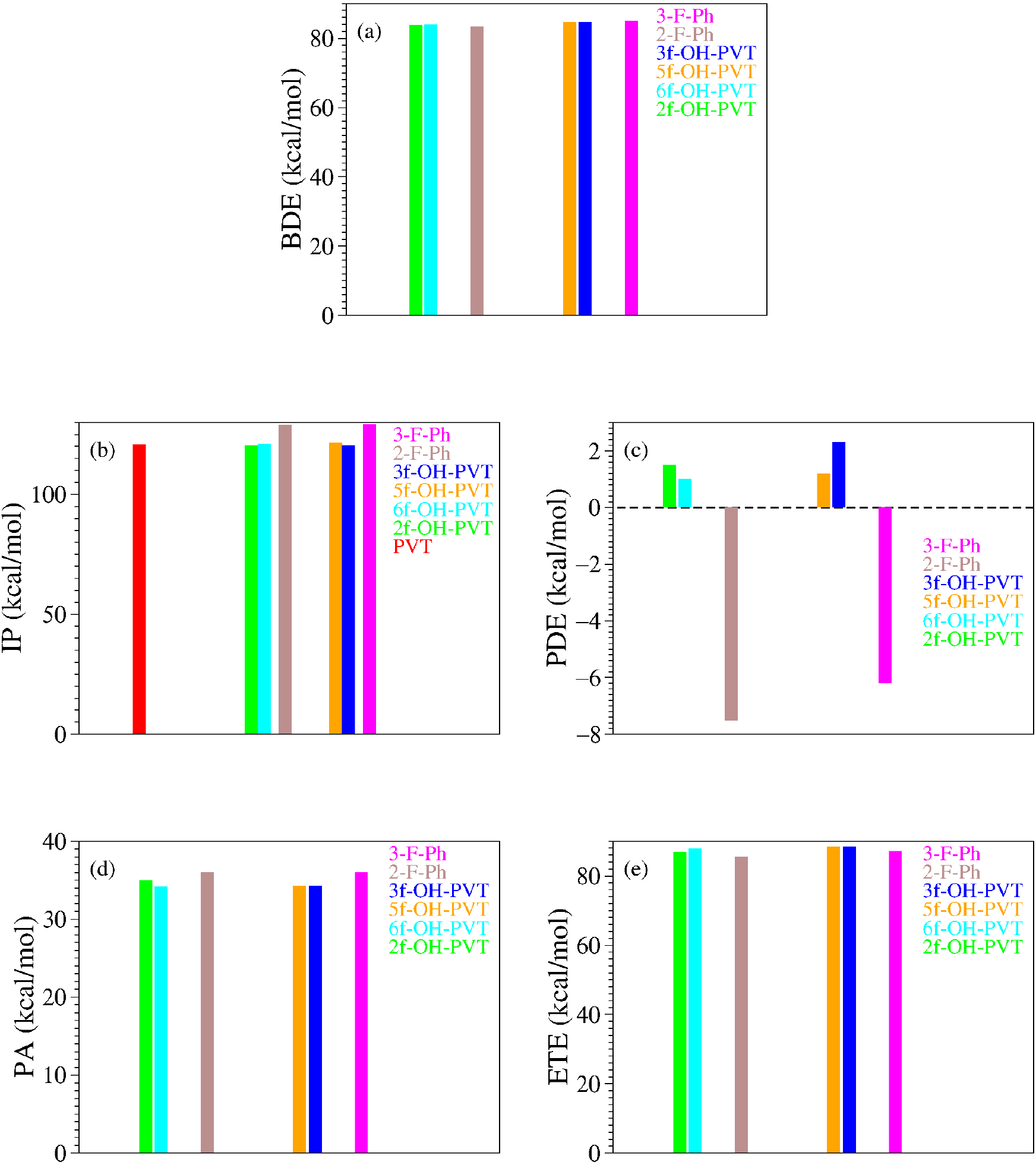
Enthalpies of reaction quantifying the antioxidant potency of F-hydroxylated PVT derivatives at positions where an H atom on the PVT’s fluorophenyl moiety was replaced by an OH group (nf-OH-PVT, *n* = 2, 3, 5, 6) depicted along with those for the isolated fluorophenol molecule corresponding to them (n-F-Ph, *n* = 2, 3): (a) bond dissociation enthalpy (BDE), (b) ionization potential (IP), (c) proton dissociation enthalpy (PDE), (d) proton affinity (PA), and (e) electron transfer enthalpy (ETE).

Inspection of Table 4 unravels that the reduction of BDE is the most important effect brought about by the additional OH group. The values of BDE characterizing the various F-hydoxylated derivatives do not significantly differ from each other; these differences fall within the chemical accuracy range (*∼* 1 kcal/mol). In spite of the fact that the values of BDE for the F-hydoxylated derivatives are by up to 7 kcal/mol larger than those for the Q-hydoxylated derivatives, they are nevertheless up to 16 kcal/mol smaller than the smallest BDE of the parent PVT. Although the F-hydoxylated PVT derivatives possess BDE values somewhat larger than the Q-hydoxylated PVT derivatives, their BDE’s should be comfortably larger than those of PVT to make them better antiradical agents than the parent PVT.

## 4. Conclusion

As emphasized recently [55], discriminating between different free radical scavenging pathways (in our specific case, whether HAT or SPLET prevails) cannot merely rely on the enthalpies of reaction of the antioxidant. To this aim, addition information on the free radical properties and/or the reaction kinetics is absolutely necessary. Based on the similarity of the presently investigated pitavastatin (PVT) and the recently studied atorvastatin (ATV) [53], one can nevertheless argue that HAT is the dominant mechanism by which PVT scavenges free radicals even in polar solvents like the presently considered methanol; if SPLET were the dominant mechanism, hdroxylated PVT metabolites (let them be of Q-type or F-type) would not be better free radical scavengers than the parent PVT but the contrary turned out to be the case of the related atorvastatin [47] and fluvastatin [58].

In this vein, we believe that the extensive results quantifying the antiradical activity of PVT and of its hydroxylated metabolites presented in this paper are quite important. Reporting that the hydroxylated PVT metabolites possess bond dissociation enthalpies (BDE, i.e., the property related to HAT) substantially smaller than the parent PVT, the present work indicates that the hydroxylated PVT metabolites are better antiradical agents than the parent PVT drug.

Noteworthily, this conclusion applies both to the (Q-)hydroxylated PVT derivatives obtained by replacement of a hydrogen atom on the quinoline core by an OH group and to the (F-)hydroxylated PVT derivatives obtained by replacement of a hydrogen atom on the fluorophenyl ring by an OH group. This is important because the (Q-)hydroxylated PVT derivatives and the (F-)hydroxylated PVT derivatives significantly differ from each other. Indeed, inspection of the spatial densities of PVT’s frontier molecular orbitals FMO (Figures 1) reveals that the former correspond to part of the PVT molecule of highest chemical reactivity. This is in contrast to the latter wherein the contribution to LUMO is altogether negligible and the contribution to HOMO is very modest. Is the change in FMO spatial distributions brought about by the additional OH group of the (Q-)hydroxylated PVT derivatives beneficial or rather detrimental for the overall effect on LDL concentration? If the latter applies, (F-)hydroxylated PVT derivatives are to be preferred notwithstanding the fact that their BDE of the extra OH group is somewhat larger than that of the (Q-)hydroxylated PVT derivatives. This is an important aspect that remains to be interrogated in subsequent in vitro and in vivo experimental investigations that the present theoretical study aims at stimulating.

## Data Availability Statement

The data that support the findings of this study are available from the author upon reasonable request.

